# Social valence dictates sex differences in identity recognition

**DOI:** 10.1101/2024.06.07.598039

**Authors:** Amanda Larosa, Qi Wei Xu, Alice S. Wong, J. Quinn Lee, Mark P. Brandon, Tak Pan Wong

## Abstract

Social valence is the directional emotional significance affiliated with social experiences. Maladaptive processing of negative social valence (NSV) has been linked to mood disorder susceptibility, which is more prevalent in women. To determine whether there are sex differences in NSV processing, we developed social valence tasks where the identity recognition of conspecifics with distinct valences served as the readout. Male mice demonstrated identity recognition regardless of social valence. Conversely, female mice did not show identity recognition following the NSV task. *In vivo* calcium imaging of the dorsal CA1 further revealed sex differences in NSV processing with reduced hippocampal representation of social information in female mice. These results suggest the imprecise encoding of NSV may contribute to the heightened vulnerability to social stress-related mood disorders in women.

## Introduction

Social relationship dynamics are an influential factor on mental health and mood disorder vulnerability (*1, 2*). Social interactions often carry emotional meaning, where negative experiences lead to behaviors that aim to prevent conflict and playful or nurturing interactions are generally rewarding. The emotional affect attached to interactions is termed social valence. Maladaptive negative social valence processing has been implicated in enhancing mood disorder susceptibility (*3*), such as in the higher vulnerability to depression in women (*4-6*). Hippocampal activity was shown to be modulated by negative (*7*) and positive socially valenced experiences (*8*). We hypothesized that sex differences in negative social valence processing could be related to disrupted hippocampal activity. Using novel behavioral assays for positive (PSV) and negative social valence (NSV) association with conspecifics, we demonstrated comparable identity recognition informed by PSV in both sexes. However, female mice, but not male mice, failed to differentiate the identity of conspecifics that were associated with NSV. We further showed that female mice displayed decreased processing of negative social information by dorsal CA1 (dCA1) neurons. Our results suggest a hippocampal mechanism for sex differences in NSV processing, which may contribute to the greater mood disorder susceptibility among women.

## Results

### Female mice demonstrate identity recognition in positive, not negative, social valence tasks

We designed behavioral tasks based on the recognition of individual conspecific identity. In PSV experiments, male and female mice were allowed to interact with two sex-matched CD1 conspecifics: a neutral mouse and a reward-associated mouse (PSV mouse) **(Fig. 1A)**. Upon exploration of the PSV mouse, a food pellet was delivered. No food reward was given during the experience with the neutral mouse. Since subject mice became familiar with both the neutral and the PSV mouse, the decision to interact in future encounters was not determined by social novelty, but instead the emotional valence associated with the identity of the conspecific mouse (e.g., spending more time with the PSV mouse that predicts reward than the neutral mouse). As we expected, one day after training in a social discrimination test (SDT) with both the neutral and PSV mice present in wire cups, both sexes spent more time exploring the PSV mouse as measured by interaction time in the SDT and normalized intersession ratios (i.e., social interaction time normalized by empty cup interaction time) **(Fig. 1A)**.

**Fig. 1:**
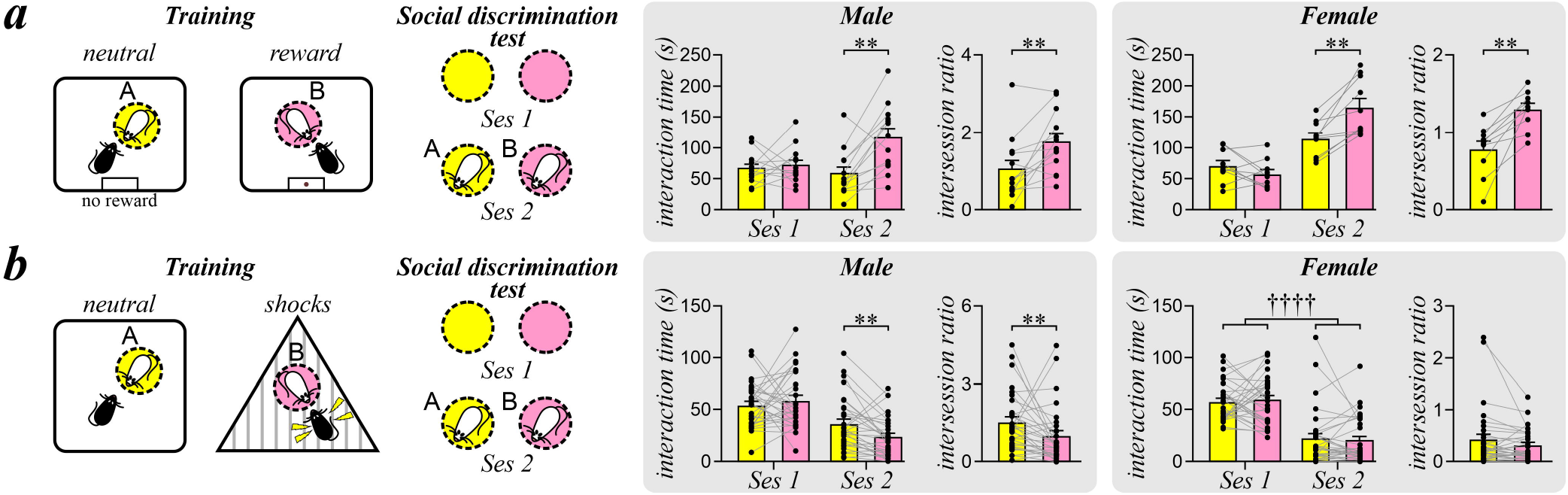
Sex difference in identity recognition in the negative social valence (NSV) task, but not the positive social valence (PSV) task. (**A**) Behavioral schematic of the PSV task. Neutral trials with mouse A (yellow) and PSV trials with mouse B (pink). Histograms show the interaction time during session 1 (Ses 1, empty cups) and session 2 (Ses 2, with conspecific mice) of the social discrimination test, and the intersession ratio (Ses 2/Ses 1 interaction time) for each conspecific. (**B**) Behavioral schematic of the NSV task. Neutral trials with conspecific A (yellow) and NSV trials with conspecific B (pink). Histograms show the interaction time and intersession ratio during the social discrimination test for each conspecific. Data are presented as mean ± SEM. ††††p < 0.0001, effect test of session with two-way ANOVA. **p < 0.01, Wilcoxon test with Bonferroni correction.

In NSV experiments we used another cohort of mice that became familiar with a neutral and a shock-associated CD1 mouse (NSV mouse). During exploration of the latter, aversive foot shocks were delivered to subject mice **(Fig. 1B)**. We expected subjects to avoid the NSV mouse in the SDT. As anticipated, male mice showed lower interaction time and intersession ratio with the NSV mouse, compared to the neutral mouse. Since neutral and NSV trials were alternated across 3 days, we reversed the trial order to address potential effects of order upon performance in the SDT and found that decreased interaction and intersession ratio with the NSV mouse were maintained in male mice **(fig. S1)**. Conversely, avoidance of the NSV mouse was not observed in female mice, who instead showed decreased interaction time with both conspecifics along with no differences in intersession ratio **(Fig. 1B)**. These findings suggest that female mice did not differentiate between the identities of these conspecifics, developing social avoidance to both NSV and neutral mice.

Sex differences in identity recognition in the NSV task were not due to diminished social memory capability. Not only did both sexes display intact identity recognition in the PSV task **(Fig. 1A)**, but they also showed a preference toward novel conspecifics regardless of their strain in social novelty tests **(fig. S2)**. Moreover, we performed the NSV task with same-strain conspecific C57BL/6 targets, which were suggested to be easier to recognize than mice from a different strain (*9*). We found that male mice, but not female mice, continued to show decreased interaction and intersession ratio with the NSV mouse **(fig. S3)**. Avoidance was not found in male and female control mice that were trained without foot shocks, **(fig. S4)**, eliminating potential preferences in conspecifics that were not dictated by prior aversive experience.

In attempts to enhance identity recognition of female mice in the NSV task, we employed several variations of the task. We acknowledged that female mice may show stronger social memory for opposite sex conspecifics (*10*) and tested the impact of using male conspecifics. Although interaction times with both conspecific mice were increased, we did not observe avoidance of the NSV mouse **(fig. S5)**. Next, we attempted to strengthen the task by delivering foot shocks following each bout of interaction with the NSV mouse in several variations of the task. First, we used 2 contexts like in **Fig. 1B** for neutral and NSV trials **(fig. S6a)**. Second, subjects were introduced to a shock box containing both conspecifics and an empty cup but were only shocked upon interaction with the NSV mouse in one **(fig. S6b)** or three training sessions **(fig. S6c)**. Lastly, female mice were trained twice in a shock box containing both conspecifics, but several contextual cues were changed between sessions (i.e., wall color, cup color and shape, training room) to reduce the influence of contextual information from the apparatus **(fig. S6d)**. In all cases, we failed to elicit a decrease in interaction or intersession ratio with the NSV mouse in female mice. Finally, since the estrous status of female mice has been shown to affect social behaviors (*11*), we separately examined the performance of sexually receptive (estrus/proestrus) and nonreceptive (diestrus/metestrus) female mice in the NSV task. Nonetheless, both groups displayed no identity recognition **(fig. S7)**. Together these findings demonstrate that negative, but not positive, social valence hinders identity recognition in female mice.

### Both sexes demonstrate recognition in positive and negative object valence tasks

To determine whether the sex difference in NSV identity recognition is due to deficits in learning to associate a stimulus with negative valence, we conducted positive (POV) and negative object valence (NOV) tasks. The POV and NOV tasks were similar to social valence experiments, but instead associated inanimate objects with either appetitive reward or foot shocks. Both male and female mice not only demonstrated increased investigation of a POV target **(Fig. 2A)**, but now showed decreased investigation of a NOV target **(Fig. 2B)**. These results were supported by parallel changes in intersession ratio. The maintenance of object recognition regardless of valence supports the notion that associative learning of negative stimuli is intact in both sexes. Preferences in investigation were influenced by object valence as neither sex demonstrated changes in exploration when both objects were neutral **(fig. S8)**. Thus, the sex difference in identity recognition is governed by the social nature of the NSV task in female mice.

**Fig. 2:**
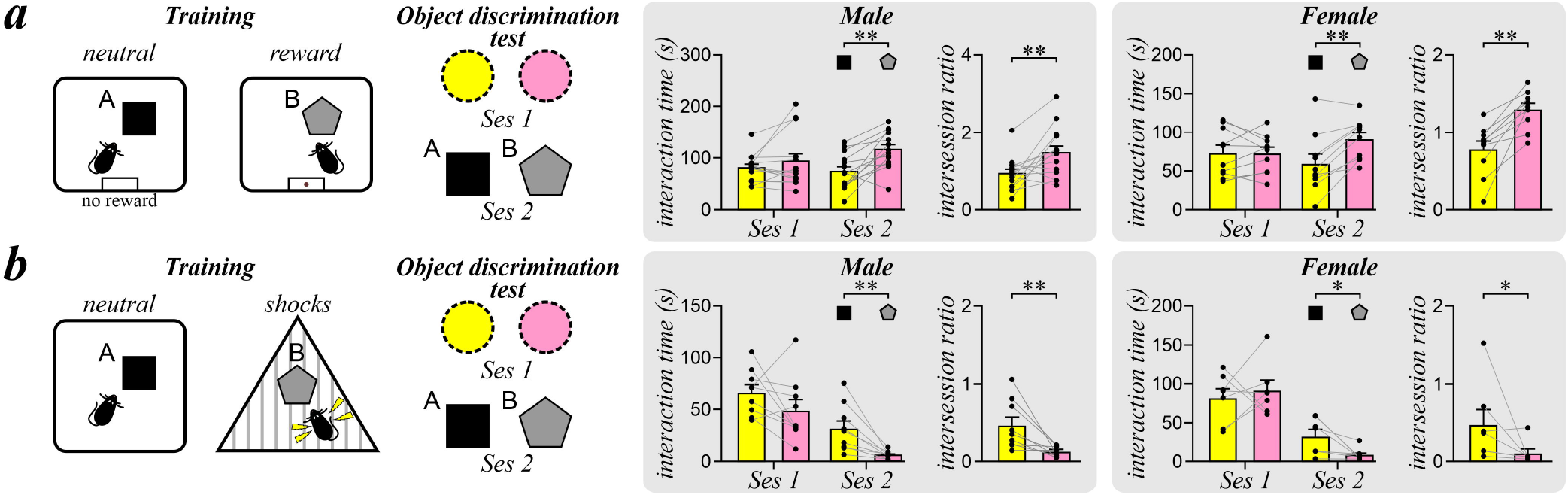
Object recognition in both sexes in the positive object valence (POV) and negative object valence (NOV) tasks. (**A**) Behavioral schematic of the POV task. Neutral trials with object A (black square) and POV trials with object B (gray pentagon). Histograms show the interaction time during session 1 (Ses 1, empty cups) and session 2 (Ses 2, objects) of the object discrimination test, and the intersession ratio (Ses 2/Ses 1 interaction time) for each object. (**B**) Behavioral schematic of the NOV task. Neutral trials with object A (black square) and NOV trials with object B (gray pentagon). Histograms show the interaction time and intersession ratio during the object discrimination test for each object. Data are presented as mean ± SEM. *p < 0.05, **p < 0.01, Wilcoxon test with Bonferroni correction.

### Female mice do not demonstrate identity recognition in a negative social valence task using social defeat

Rodents are territorial animals. Aggressive encounters between conspecifics are important for establishing social hierarchy and have been previously used to elicit identity recognition in hamsters (*12*). Hamsters were trained to become familiar with two aggressive hamsters, one through a partition (neutral) and the other with direct contact that allowed for physical attacks (agonistic). Animals avoided the agonistic conspecific supporting identity recognition. With our experience in studying social defeat (*7*), we developed a NSV task using social defeat stress (NSV-SDS) by modifying the hamster model. We used two aggressive mouse strains to examine sex differences in NSV-SDS identity recognition. For male mice, we used aggressive CD1 males that are commonly used in chronic social defeat stress. Since female CD1 mice are not aggressive, we adopted a female rival aggression model to identify aggressive Swiss Webster females by housing them with a castrated male (*13*). During training in the social defeat task, subjects were exposed to a neutral and an NSV-SDS mouse who they experienced social defeat attacks from. To eliminate potential pheromonal differences, both neutral and NSV-SDS mice were aggressors, but the former was confined to an enclosure and thus could not attack, while the latter was freely moving. One day after training in the SDT, male mice spent less time interacting with the NSV-SDS mouse and showed decreased intersession ratios compared to the neutral mouse **(Fig. 3)**. However, female subjects continued to show no difference in interaction time or intersession ratio between conspecifics.

**Fig. 3:**
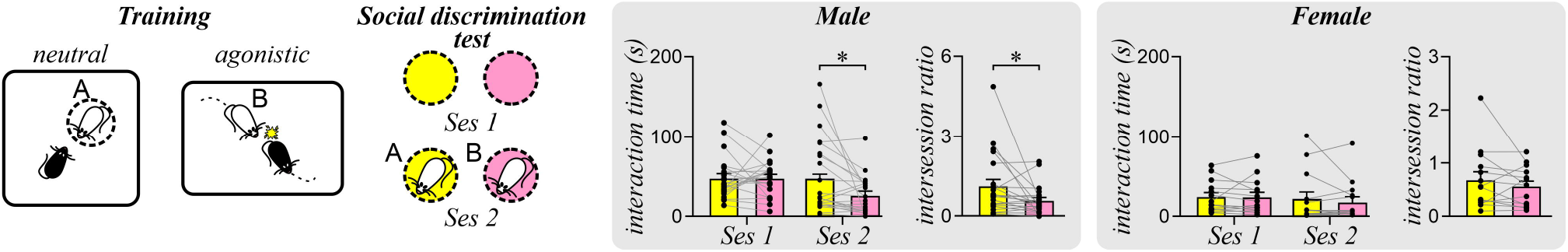
Sex difference in identity recognition in the negative social valence-social defeat stress (NSV-SDS) task. Behavioral schematic of the NSV-SDS task. Neutral trials with mouse A (yellow) and NSV-SDS trials with mouse B (pink). Histograms show the interaction time during session 1 (Ses 1, empty cups) and session 2 (Ses 2, with conspecific mice) of the social discrimination test, and the intersession ratio (Ses 2/Ses 1 interaction time) for each conspecific. Data are presented as mean ± SEM. *p < 0.05, Wilcoxon test with Bonferroni correction.

Several experiments were conducted to confirm that male mice discriminated between conspecifics through a learned understanding of both individuals. First, we trained subjects with the neutral and NSV-SDS mice but conducted testing with the neutral and a novel mouse (**fig. S9a)**. We found that subjects had increased interaction time and intersession ratio with the novel mouse, indicating that they recognized the neutral conspecific as familiar. These findings suggest that the avoidance of the NSV-SDS mouse in the previous experiment **(Fig. 3A)** was not due to a lack of familiarity with the neutral mouse. Second, to determine whether decreased interaction with the NSV-SDS mouse was governed by the aversive social experience, we trained male mice with the neutral and NSV-SDS mice but tested them with two novel conspecifics. No change in interaction or intersession ratio was observed with either conspecific **(fig. S9b)**. Moreover, males that interacted with both conspecifics in the neutral and SDS contexts, but in the absence of social defeats did not show differences in interaction or intersession ratio **(fig. S9c)**. Together these findings indicate that male mice recognize both the neutral and NSV-SDS mice as familiar and only attribute the aversive experience with the latter.

We acknowledged that interactions with the partitioned neutral mouse may differ in somatosensory quality from the direct attacks experienced with the NSV-SDS mouse. For example, the opportunity for anogenital sniffing and direct physical touch with the NSV-SDS mouse is diminished with the neutral mouse that was confined to an enclosure. However, identity recognition was not rescued in female mice even when we allowed freely moving interactions of female subject mice with neutral nonaggressive Swiss Webster mice that did not attack **(fig. S10a)**. Lastly, control female mice who interacted with both conspecifics in the absence of social defeat stress showed no change in interaction or intersession ratio during the SDT **(fig. S10b)**. It is possible that the lack of avoidance of the NSV-SDS may be in part attributed to a fewer number of attacks from female compared to male aggressors **(fig. S11a)**. However, attack number did not correlate with SDT interaction time with either conspecific in both males and females, indicating that differences in identity recognition cannot be explained by the number of agonistic encounters **(fig. S11b)**. Taken together, we concluded that female mice show diminished NSV identity recognition compared to male mice.

### Female mice show a decrease in dCA1 correlates of social interaction following a negative social valence task

The hippocampus is a region known to support social cognition (*14*), with proper function of the dorsal CA2 (*15*), dCA1 (*12*), and ventral CA1 (*16*) subregions required for social memory. The dCA1 is also necessary for successful identity recognition following valenced social learning (*12, 17*). To determine whether the dCA1 is required for identity recognition in the NSV-SDS task, we inhibited N-methyl-D-aspartate receptor (NMDAR) activity in the dCA1 that supports synaptic plasticity mechanisms underlying social memory functions (*18, 19*). We found that blocking NMDARs in the dCA1 by bilateral infusions of the NMDAR antagonist APV prior to the neutral and NSV-SDS training trials abolished identity recognition. However, vehicle-infused subjects maintained avoidance of the NSV-SDS mouse **(fig. S12)**. Although central to social memory mechanisms (*15*), the adjacent CA2 subregion appears resistant to NMDAR-mediated plasticity (*20*) and plasticity mechanisms associated with social memory in the CA2 are NMDAR-independent (*18*). Thus, dCA1 functioning is required for identity recognition in the NSV-SDS task.

More recently, dCA1 population activity was shown to be selective for individual social targets in male C57BL/6 mice (*17*). Since hippocampal activity is highly sensitive to stress (*21*), we hypothesized that there are sex differences in social information representation in the dCA1 in a stressful task like NSV-SDS. We conducted *in vivo* calcium imaging in dCA1 of freely-behaving mice during the SDT that followed the NSV-SDS task **(Fig. 4, A and B)**. Comparing the mean inferred rate of calcium spikes from all dCA1 neurons during the 2 sessions of the SDT (session 1: two empty wire cups; session 2 neutral and NSV-SDS mice in each of these cups), revealed lower spiking rate in female mice than male mice during both SDT sessions. In addition, we found that dCA1 neurons fired less during session 2 than session 1 in female mice only.

**Fig. 4:**
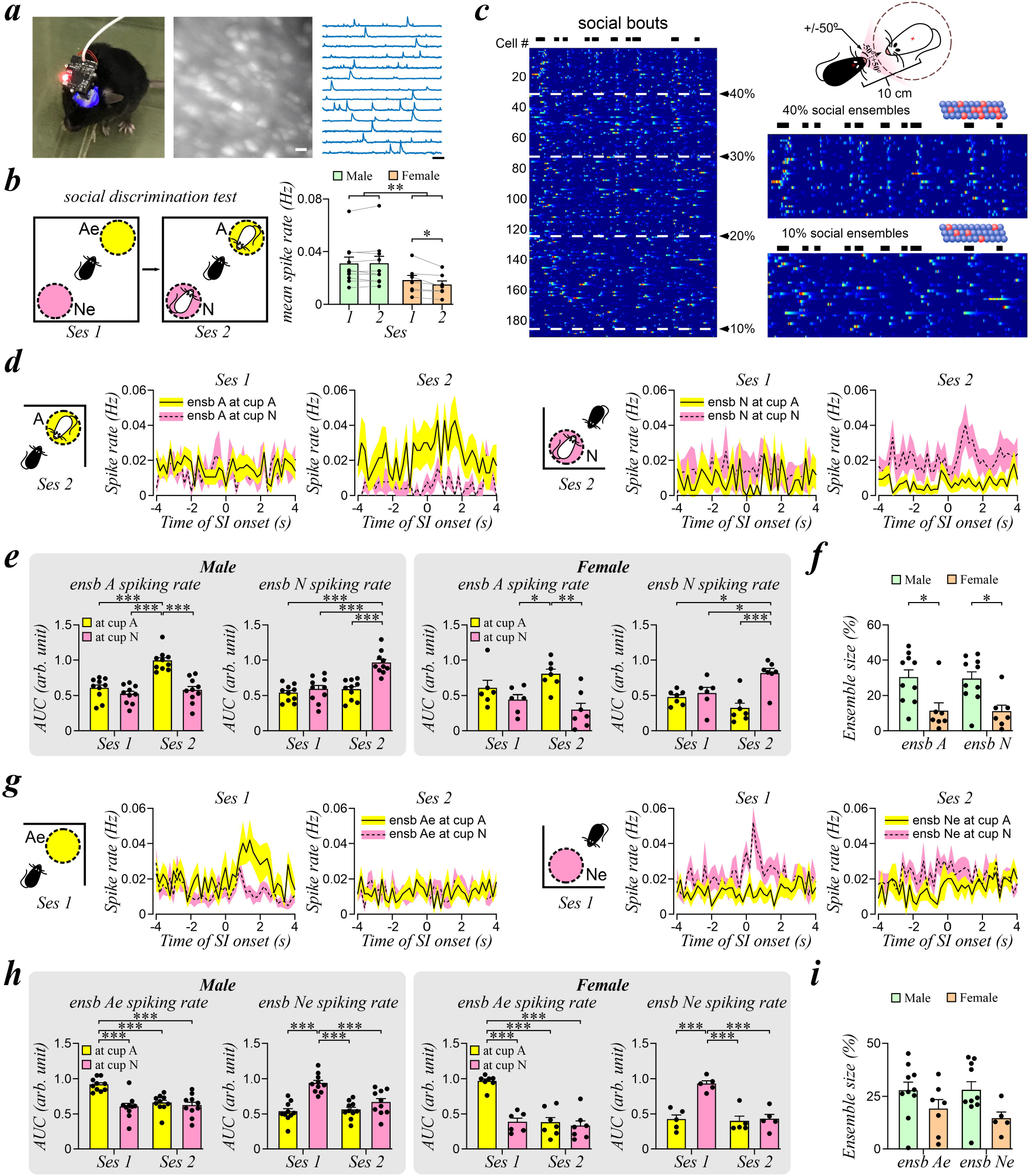
Sex differences in dCA1 neuronal activity during identity recognition. (**A**) *Left:* A subject mouse with a miniscope for *in vivo* calcium imaging. *Middle:* Micrograph from *in vivo* calcium imaging recording showing individual GCaMP6f-expressing dCA1 neurons (scale bar = 20 μm). *Right*: Traces represent GCaMP6f fluorescent signals from dCA1 neurons (scale bar: 1 min). (**B**) *Left*: Behavioral schematic of the social discrimination test (SDT). Subjects explore in session 1 (Ses 1, empty cups) and session 2 (Ses 2, with conspecific mice). Agonistic (A: yellow) and neutral (N; pink) social targets from the NSV-SDS task. *Right*: Overall firing rate of dCA1 neurons in Ses 1 and Ses 2 of the SDT in male (green) and female (orange) subjects. **p < 0.01, Effect test of sex from two-way ANOVA. *p < 0.05, Wilcoxon test with Bonferroni correction. (**C**) Social interaction was defined as < 10 cm and ± 50° between the head and heading direction of the subject mouse, respectively, and the center of a wire cup containing the target mouse. *Left:* The calcium signal for each dCA1 neuron was sorted according to their activity during bouts of social interaction (black rectangles above raster plots). *Right:* Social ensembles of dCA1 neurons that were active during 40% (top) or 10% (bottom) of social bouts. (**D**) Line plots of mean activity (± SEM) of dCA1 ensembles for mouse A (ensb A, *left*) and mouse N (ensb N, *right*) during interaction with cup A and N in different sessions of SDT. (**E)** Histograms of the area under curves (AUC) of averaged ensb A and ensb N activity during interactions with different cups in *Ses 1* and *Ses 2* in male (*Left*) or female subjects (*Right*). *p < 0.05, **p < 0.01, *** p < 0.001, Wilcoxon test. (**F**) Ensemble size for ensb A or ensb N in male and female subjects. *p < 0.05, Student’s t-test. (**G**) Line plots of mean activity (± SEM) of dCA1 ensembles for empty cup A (ensb Ae, *left*) and empty cup N (ensb Ne, *right*) during interaction with cup A and N in different sessions of SDT. (**H**) Histograms of the area under curves (AUC) of averaged ensb Ae and ensb Ne activity during interactions with different cups in *Ses 1* and *Ses 2* in male (*Left*) or female subjects (*Right*). *** p < 0.001, Wilcoxon test. **(I)** Ensemble size for empty cups Ae or Ne in male and female mice. Data are presented as mean ± SEM.

We examined dCA1 population activity that is related to social interaction in the SDT. We focused on neurons that were active during at least 40% of all social interaction bouts in session 2 of the SDT **(Fig. 4C)**. The mean activity vector of these neurons and the social interaction bout vector displayed a significant cosine similarity index (CSI; (*22*)) when compared to shuffled social interaction bout vector or other behavioral vectors (e.g., speed, head direction, empty cup, or another social target). We named them social ensembles and isolated social ensembles for the neutral mouse (ensemble N) and the agonistic NSV-SDS mouse (ensemble A). The activity of a social ensemble was specific to its associated conspecific target. As shown in **Fig. 4D**, the social ensemble A activity was elevated during interactions with the agonistic NSV-SDS mouse and showed lower activity during interactions with the neutral mouse or with the empty cups in session 1 of the SDT **(Fig. 4E)**. Social ensemble N activity was similarly selective for interactions with the neutral mouse in session 2 of the SDT. Since the cup location between SDT sessions was unchanged, the elevations in ensemble activity in session 2 only were presumably due to conspecific presence and not visiting the same spatial location.

Next, we compared social ensemble activity between male and female mice. In male mice, the activity of social ensembles A and N were highest during interactions with the agonistic NSV-SDS mouse and neutral mouse, respectively, supporting a high specificity. However, in female mice, social ensemble A activity during interactions with the agonistic NSV-SDS mouse were similar to those during interaction with that same empty cup in session 1. In other words, even though a conspecific was introduced to the cup, social ensemble activity did not reflect this change, which may suggest a loss in specificity to represent interactions. Since social ensembles are sparsely distributed, we compared the size of these social ensembles (number of dCA1 neurons in social ensembles / number of all recorded dCA1 neurons) and found a significant decrease in both ensemble A and N size in female compared to male subjects **(Fig. 4F)**.

Although the decreased specificity and size of dCA1 social ensembles in female mice is in line with the sex differences in identity recognition, these properties may result from an overall lower activity of dCA1 neurons **(Fig. 4B)**. To determine whether the reduced dCA1 ensemble properties in female mice can also be found during interactions with nonsocial objects, we identified dCA1 nonsocial ensembles that are specific for the empty wire cup A or cup N in session 1 of the SDT. Like social ensembles, nonsocial dCA1 ensembles exhibit higher activity during visits to the corresponding empty cup **(Fig. 4G)**. However, the nonsocial ensembles of both male and female mice displayed high specificity **(Fig. 4H)**. In addition, no differences between ensemble size of nonsocial ensembles were found between male and female mice **(Fig. 4I)**. Finally, when we compared the CSI of social and nonsocial ensemble vectors with the corresponding social target or object vectors between male and female mice, we found a significant decrease in CSI of social ensembles in female mice only **(fig. S13)**. Taken together, compared to male mice, we found that female mice exhibited a decrease in specificity and size of dCA1 social ensembles, but not nonsocial ensembles.

## Discussion

Our data demonstrate that both male and female mice can identify individuals, while female mice cannot identify a conspecific that is associated with a NSV. Importantly, this observed sex difference cannot be attributed to impaired social memory capacity in female mice, as evidenced by sustained PSV identity recognition **(Fig. 1A)** and social novelty preference **(fig. S2)**. This also does not stem from deficits in associative learning, as in addition to PSV, female mice exhibit valenced learning of object identity in the POV and NOV tasks **(Fig. 2)**. Despite attempts at enhancing recognition by optimizing task designs, such as utilizing same strain **(fig. S3)** or opposite sex NSV mice **(fig. S5)**, manipulating temporal associations in NSV **(fig. S6)**, and equalizing somatosensory quality in NSV-SDS **(fig. S10)** no improvement in identity recognition was observed in female mice. Moreover, hormonal influence from differences in estrous cycle stage do not account for these effects **(fig. S7)**. Our findings therefore support a specific effect of NSV on identity recognition unique to female mice.

The dCA1 subregion of the hippocampus is thought to encode spatial information pertaining to other conspecifics in the environment (*23, 24*), representing a critical aspect of social cognition. Additionally, dCA1 is required for valenced identity recognition (*12, 17*). We showed that the dCA1 is required for NSV-SDS in male subjects and that neurons were less reliable in their representation of social interactions in female compared to male mice **(fig. s12, Fig. 4)**. We propose that this reduced fidelity of the dCA1 in the representation of interactions may underlie the diminished NSV identity recognition in female mice. This decreased reliability was specific to social ensembles, but not nonsocial ensembles, mirroring the accurate identification of objects, but not conspecifics.

Our findings that female subjects avoided both the neutral and NSV mouse in the SDT suggest a generalization of negative memories, a shared cognitive symptom of depression, post-traumatic stress disorder, and anxiety which are more prevalent in women (*6, 25-28*). Fear generalization to cues that do not predict threat may be promoted by elevations in hippocampal corticosterone (*29, 30*) and accompanied by altered hippocampal function in female rodents (*31, 32*). Contextual fear generalization can also be induced by NSV and is mediated by extended circuitry to and from the dorsal hippocampus (*33*). Previous findings using the same NSV social defeat model in female mice suggested that resulting hypervigilance, demonstrated as an exaggeration in defensive behaviors towards neutral mice, may underlie the inability to discriminate threatening from non-threatening social targets (*13*). Thus, the less stressful nature of the social novelty test and PSV task may explain why female mice are able to demonstrate identity recognition.

Our findings have implications on NSV processing in humans. While women typically outperform men in facial recognition tasks (*34, 35*), they exhibit more errors than male participants when negatively valenced information is introduced (*36*). Women spend more time processing negative faces (*37*) and show greater limbic (*38, 39*) and cortisol responses (*5*) compared to men. This supports our hypothesis that NSV imparts greater consequences in women. There remains limited knowledge regarding the involvement of hippocampal processing in valenced social interactions, but some clinical support exists. Hippocampal connectivity with limbic regions is modulated by acute social stressors (*40*) and activity increases in the learned decision to approach or avoid valenced stimuli (*41*). The hippocampus also emerges as a region of interest in studies examining brain regions differentially affected by negatively valenced stimuli between the sexes (*42, 43*). Collectively, our findings offer insights into the impact of social valence on hippocampal function between the sexes, with potential implications for understanding mood disorder susceptibility and social cognition.

## Acknowledgments

We would like to thank the UCLA miniscope team and Daniel Aharoni’s lab for developing and sharing the miniscope technology. The present study used the services of the Molecular and Cellular Microscopy Platform in the Douglas Hospital Research Centre. Melina Jaramillo Garcia helped set up the imaging verification. We also thank Dr. Rosemary Bagot for the helpful discussions and advice.

## Funding

Natural Sciences and Engineering Research Council of Canada 8401073 (TPW)

Canadian Institutes of Health Research PJ8 179866 (TPW)

Fonds de recherche du Québec – Nature et technologies 326838 (TPW)

## Author contributions

Conceptualization: AL, QWX, MPB, TPW

Methodology: AL, QWX, ASW, JQL

Investigation: AL, QWX

Funding acquisition: TPW

Project administration: ASW, TPW

Supervision: TPW

Writing – original draft: AL, TPW

Writing – review & editing: AL, QWX, JQL, MPB, TPW

## Data and materials availability

All data are available in the main text or the supplementary materials. Data and codes are available upon request.

## Supplementary Materials

Materials and Methods

Supplementary Text

Figs. S1 to S13

References (*1–43*)

