## Supplementary material for "Social valence dictates sex differences in identity recognition": Methods and supplementary figures

**The PDF file includes:**

Materials and Methods
Figs. S1 to S13

### Materials and Methods

Animals

Male and female adult C57BL/6 mice (8-12 weeks old), CD1 retired breeders (3-6 months-old), and Swiss Webster resident pairs (3-6 months-old) were obtained from Charles River. C57BL/6 mice to be used as conspecific social targets were obtained from Jackson Laboratory to ensure they would be from different litters as the subject mice. In all experiments, sex-matched conspecifics were used to eliminate preferences towards the opposite sex, unless otherwise specified. All experiments were approved by the Facility Animal Care Committee at the Douglas Institute and followed guidelines from the Canadian Council on Animal Care (protocol no.: DOUG-5935).

Positive social valence (PSV) and positive object valence (POV) tasks

To determine whether an individual conspecific or inanimate object could be associated with a positively-valenced experience we developed appetitive PSV and POV tasks. *Food deprivation:* Subjects were food restricted for 6-8 hours prior to habituation to encourage the consumption of food rewards. Animals typically lost 1-1.5 g, translating to a loss of less than 10% of their body weight. *Habituation:* Mice were habituated to the food delivery process for 3 days where they were allowed to explore a modified empty rat cage containing a food dish below a delivery tube. A chocolate food pellet (20 mg, BioServ) was manually delivered every minute and the number of pellets consumed within 10 minutes was recorded. Animals that failed to consume any food pellets were excluded from further training (n = 1). *Training:* Training was conducted over two consecutive days where subjects were exposed to a reward (PSV or POV) and a neutral trial. The inter-trial interval was 2 hours and the trial order was counterbalanced. Both trials took place in a rectangular cage (25 cm x 37 cm) with white walls and a paper floor. During the PSV trial, subjects freely explored a CD1 mouse (PSV mouse) in a perforated enclosure adjacent to the food dish for 10 minutes. Upon each bout of interaction with the PSV mouse, subjects received one food pellet. The number of interactions and pellets consumed were recorded. In the neutral trial, a different CD1 mouse was placed in the same location, but no food reward was given upon interactions. PSV and neutral mice were similar in age and size (± 5 g). During the inter-trial interval, subjects were returned to their homecages. We also conducted POV training in another mouse cohort by replacing conspecifics with inanimate objects. These objects were dissimilar in appearance and touch (i.e. silicone vs. plastic; dark vs. light in color; circular vs. irregular shape), but similar in size. Objects were randomly assigned to either POV or neutral trials. All behavioral experiments were performed under ambient red light, with static white noise at 60 dB.

Negative social valence (NSV) and negative object valence (NOV) tasks

To determine whether an individual mouse could be associated with a negatively-valenced experience we developed several negative valence tasks using electric foot shocks (NSV) or social defeat stress (NSV-SDS). In addition, a NOV task was developed to associate electric foot shocks with inanimate objects.

NSV

*Habituation:* Two hours prior to the first training session, mice were habituated to two contexts for 5 minutes. NSV trials took place in a triangular shock box (46 cm x 46 cm x 46 cm) with black walls and a metal grid floor, while neutral trials were conducted in a rectangular cage (25 cm x 37 cm) with transparent walls and a paper floor. Subjects explored each context which contained empty perforated enclosures (10 cm x 5 cm x 30 cm) at the centre. This allowed subjects to be familiarized with both contexts, encouraging interaction with conspecifics during training, rather than context exploration. *Training:* Training was conducted over 3 consecutive days where subjects were exposed to a NSV and a neutral trial. Each trial was 5 minutes long, with an inter-trial interval of 2 hours. The trial order was counterbalanced. In the NSV trial, foot shocks (0.3 mA, 1 s) were delivered at 4 and 4.5 minutes, to allow sufficient interaction time with the NSV mouse prior to shock delivery. An insulating tape coated bottom was placed under the enclosure to protect the NSV mouse from foot shocks. In the neutral trial, no foot shocks were given. During the inter-trial interval, subjects were returned to their homecages. The conspecifics were CD1 or C57BL/6 mice as specified in the results.

NSV and NOV (interaction dependent)

We attempted to enhance the association between the negative experience and conspecific identity or objects by delivering foot shocks following each interaction bout. The habituation procedure remains the same. Training procedures in different experiments include i) one 5-minute trial per day for 1 day; i) one 5-minute trial per day for 3 days; ii) three consecutive 3-minute trial per day for 1 day (used in NOV). The interval between NSV/NOV and neutral trials was always 2 hours.

NSV-Social defeat stress (NSV-SDS) task

*NSV-SDS mice:* For male experiments, aggressive CD1 retired breeders, which are commonly used in chronic social defeat stress studies, were used. Aggression towards female mice does not naturally occur in laboratory settings. To study identity recognition in female subjects, we adapted a social defeat paradigm using an ethologically valid model of rodent female aggression (*1*). In this model, aggression towards female mice is elicited by cohousing Swiss Webster females with castrated same-strain males. When the male partner is replaced for an intruder female, Swiss Webster females demonstrate rival aggression to attack the intruder.

*Training:* The NSV-SDS trial took place in a rectangular cage (25 cm x 45 cm) containing a freely moving NSV-SDS mouse and the neutral trial was conducted in an open field (44 cm x 44 cm) with a perforated enclosure at the centre, which contained a different mouse of the same strain and aggressiveness as the NSV-SDS mouse. During training, subjects explored each context during 3 trials for 3 minutes, with a 3-minute intertrial interval. In female subject experiments, the negative context was the homecage of the Swiss Webster pairs, which contained olfactory cues from the resident male and female conspecifics that may facilitate the subsequent identification of the NSV-SDS mouse. To make the olfactory cues in both interactions comparable, bedding from the homecage that the neutral female shared with their male partner was placed in the neutral context. The mouse in the neutral trial was also an aggressive Swiss Webster female. Once a trial was completed, subjects were returned to their homecages for 2 hours before beginning the other trial.

Trials with the NSV-SDS mouse were ended early if the subject jumped out of the cage to escape or showed a submissive posture (i.e., on their back). Subjects (n = 5) were removed if in the 3 attack trials, they experienced less than 15 total attacks and at least 1 non-escape trial with no attacks. Unless otherwise specified, all male CD1 and female Swiss Webster NSV-SDS mice, including neutral and other animals used in control experiments, were aggressive mice. This eliminated potential effects of differential pheromonal cues between aggressive and nonaggressive conspecifics. Although neutral conspecifics were also aggressors, they were prevented from attacking subjects as they were constrained within enclosures during interactions.

Discrimination tests

Discrimination tests were used to quantify investigation time of conspecifics and inanimate objects associated with different valences. A three chambered arena (each chamber measured 23 cm x 28 cm; 8-minute sessions) was used for all positive and negative valence experiments using foot shocks. An open field (44 cm x 44 cm; 5-minute sessions) was used for all NSV-SDS experiments. In the first session, the testing arena contained 2 empty wire cups. In the second session, the cups contained familiar mice or objects from previous training sessions, unless otherwise stated.

Three chambers social novelty preference test

The tests consisted of three 8-minute sessions: Mice were placed in the centre chamber of a three chambered arena and allowed to explore the apparatus. In the first session, the adjacent chambers contained empty wire cups. In the second session, one of the adjacent chambers now held a cup containing a novel conspecific. In the last session, one adjacent chamber held a cup containing the now familiar conspecific from the previous session, while the cup in the opposite chamber held a novel conspecific. Mice are known to prefer interactions with a novel conspecific (*2*). Conspecific location was counterbalanced for all sessions and social targets were sex-matched C57BL/6 or CD1 mice.

Estrous monitoring

Estrous monitoring was conducted in female mice that underwent the NSV paradigm. Monitoring was conducted for a total of 7 days, beginning 3 days prior to the start of behavioral training and ending the day of the SDT. Samples were collected at approximately the same time each day in the morning, prior to behavioral training and testing. Vaginal lavage was performed by flushing 0.9% NaCl into the vaginal canal with a transfer pipette. Samples were stored in a 96-well plate and stored at 4 °C. A tissue culture microscope (Nikon Diaphot) at 10x magnification was used to assess estrous cycle stage based on previously described guidelines (*3*). Results were independently verified by two experimenters and grouped into high estrous (proestrus/estrus) and low estrous (metestrus/diestrus) stages.

Surgeries

For all stereotaxic procedures, C57BL/6 mice were anesthetized with isoflurane and received subcutaneous injections of an analgesic (carprofen: 20mg/ml) during surgery and 3 days post-surgery.

*Cannula implantation*

Bilateral implantation of cannulas for drug infusion were targeted to the dCA1 of male mice (AP: -1.70; ML: ± 1.50; DV: -1.50). One week after cannula implantation, mice underwent NSV-SDS training. Bilateral intrahippocampal infusions were delivered 15 minutes prior to the first training session in each context (i.e., before the neutral and NSV-SDS trials). Subjects received an infusion of the NMDAR antagonist APV *(*1 mM, 0.5 µl*)* or vehicle (PBS). Following the termination of experiments, mice were anesthetized and perfused using PBS-heparin and 4% paraformaldehyde (PFA). Brains were extracted, fixed overnight in PFA, and cryoprotected in 30% PFA-sucrose before being frozen at -20 °C and sectioned at 45 µm thick using a cryostat

(Leica Microsystems). Sections were mounted on a slide and cannula placement was verified using a bright field microscope.

*In vivo calcium imaging*

Our surgical procedure for *in vivo* calcium imaging was previously described (*4*). Male and female C57BL/6 mice were unilaterally injected with a viral construct expressing the calcium indicator GCaMP6f (AAV2/9-SYN-GCaMP6f; 350 nL). Two sets of coordinates targeting the dCA1 were tested and achieved comparable results (set 1: AP: -1.60; ML: +1.70; DV -1.40 and set 2: AP: -1.90; ML: +1.40; DV: -1.10). One week post-injection, animals underwent a craniotomy at a 1 mm radius from the injection site. The cortical tissue above the injection site was aspirated using a blunt syringe needle connected to a vacuum pump. A gradient refractive index (GRIN) lens (0.25 pitch, 1.8 mm diameter, 4 mm length, Edmund Optics) was then lowered and secured in place with dental cement above the injection site. A metal baseplate was cemented three weeks later to allow for docking of the UCLA Miniscope (v.3). Subjects underwent NSV-SDS training. *In vivo* calcium imaging was conducted during the SDT. Following the termination of experiments, mice were anesthetized and perfused using PBS-heparin and 4% PFA. Brains were extracted, fixed overnight in PFA, and cryoprotected in 30% PFA-sucrose before being frozen at -20 °C and sectioned at 45 µm thick using a cryostat (Leica Microsystems). Sections were mounted on a slide using Fluoromount Aqueous Mounting Medium (Sigma-Aldrich). GCaMP6f expression and GRIN lens placement were verified using an automated fluorescence microscope (Olympus BX63).

Behavioral analysis

For calcium imaging experiments, behavioral analysis was performed from videos that were annotated by DeepLabCut. For other experiments, videos were analyzed manually. Interaction time was visually analyzed and included time when subjects were within 5 cm and were sniffing or oriented towards the cup. Periods where animals were <5 cm from the cup but were oriented away or grooming were not considered as social interaction. For NSV experiments, interaction time with both mice during training was measured similarly, but excluded instances where subjects were climbing the enclosure to evade foot shocks.

An intersession ratio was calculated to control for innate difference in exploratory drive and side preference. Intersession ratios were calculated for each subject by taking the ratio of the time spent interacting with a conspecific or inanimate object and the time spent interacting with an empty cup at that same location. For example: intersession ratio for target A = Session 2 interaction time with target A/Session 1 interaction time with empty cup A.

*In vivo* calcium imaging analysis

*In vivo* calcium imaging was conducted using a UCLA miniscope (v3; miniscope.org). Calcium imaging videos were recorded at 30 frames per second, together with behavioral video from a webcam at 30 frames per second using the Miniscope-DAQ-QT-Software (Aharoni-Lab, github). Concatenated video files of calcium imaging were motion corrected and aligned using Non-Rigid Motion Correction (NoRMCorre, (*5*)). Videos were downsampled to 10 frames per second and black borders that were visually deemed to have no cellular signal were cropped. Constrained Non-negative Matrix Factorization for microEndoscopic data (CNMF-E) was used to identify cell segments and extract calcium transients (*6*). Segments with spatial footprints that shared a 60% overlap and visually demonstrated comparable calcium transient dynamics were discarded as they were not considered to be individual cells. The rising phases of calcium transients were detected and binarized. They were finally downsampled to 5 Hz using customized codes in MATLAB.

*Defining social ensembles*

In behavioral videos that were recorded simultaneously during *in vivo* calcium imaging, the position and head direction of subjects in relation to the wire cups were annotated by DeepLabCut, a markerless, deep neural network-based software (*7*). Population activity of dCA1 neurons in the form of calcium signal vector of social ensembles (social ensemble vector) was calculated from the mean inferred firing rate of neurons that were active in 40% of all social interaction bouts during session 2 of the SDT. Social interactions were defined by head position and direction as timepoints when the head position of subjects was within 10 cm from the center of the wire cup, and the heading direction of subject's head was within + or – 50º (i.e., 100º). Only social interactions that lasted for at least 5 frames (i.e., 1 second) were considered social bouts. The social ensemble vector and the social interaction vector (binary logical values of social interaction frames) were used to calculate a cosine similarity index (CSI, (*8, 9*) CSI $=2B \cdot C/(|B|^2 \cdot |C|^2)$) where $C$ is the social ensemble vector (mean inferred calcium spikes) and $B$ is the behavioral vector. CSI of the social ensemble vector vs. the behavioral vector for social interactions are higher than 95% of CSIs calculated from 10,000 shuffled behavioral vectors by randomly arranging 8 equally divided portions of the behavioral vector of social interactions. We also calculated the CSI between social ensemble vectors and other behavioral vectors such as subject mouse running speed, head direction, or interaction with another mouse or empty cups and found that no significant similarities. Finally, calcium activity of nonsocial ensembles was calculated the same way as that of social ensembles, with the use of behavioral vectors from interactions with a specific empty (nonsocial) cup.

To compare sex differences in dCA1 activity, we examined the mean firing rate of all recorded neurons and neurons in social (and nonsocial) ensembles. To calculate the mean inferred spiking rate of all dCA1 neurons, we averaged the inferred calcium firing rate during different sessions of SDT from male and female mice. For the spiking rate of social and nonsocial ensembles, we averaged the inferred calcium spiking rate from dCA1 neurons that were active in at least 40% of interaction bouts with social and nonsocial targets, respectively. We also compared the size of ensembles by dividing the number of dCA1 neurons in these ensembles by the total number of recorded dCA1 neurons in each mouse.

Statistical analysis

All statistical analyses were performed using Prism 9 (GraphPad). Normality of data was determined by the Shapiro-Wilk's test. Wilcoxon test for non-parametric pairwise comparisons was used to examine interaction time with each cup within sessions. The Bonferroni correction was used to correct for multiple comparisons. Intersession ratio data was analyzed with Wilcoxon test for non-parametric pairwise comparisons. Students' t-tests were used for pairwise comparisons of parametric data. For *in vivo* calcium imaging data, two-way repeated measures ANOVA was conducted with post-hoc Tukey's test. All data were presented as mean ± SEM.

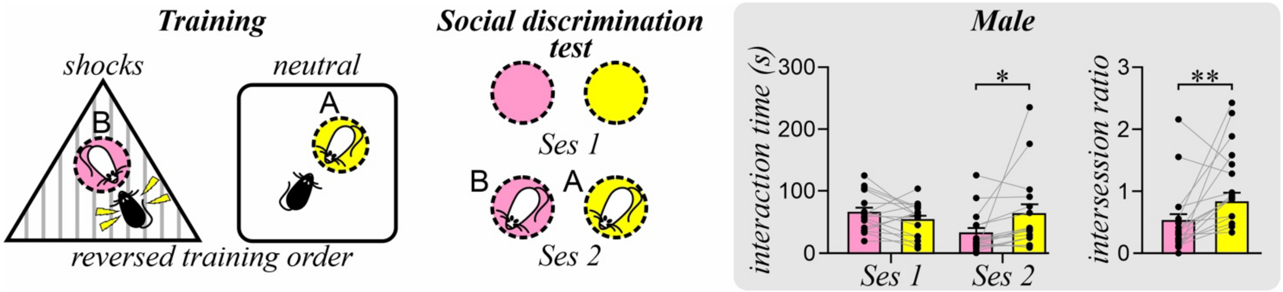


**Fig. S1. Identity recognition in male mice after the negative social valence (NSV) task with reversed training order.**

Behavioral schematic of the NSV task using a reversed training order. Neutral trials with mouse A (yellow) and NSV trials with mouse B (pink). Histograms show the interaction time during session 1 (Ses 1, empty cups) and session 2 (Ses 2, with conspecific mice) of the social discrimination test, and the intersession ratio (Ses 2/Ses 1 interaction time) for each conspecific. Data presented as mean ± SEM. *$p < 0.05$, **$p < 0.01$, Wilcoxon test with Bonferroni correction.

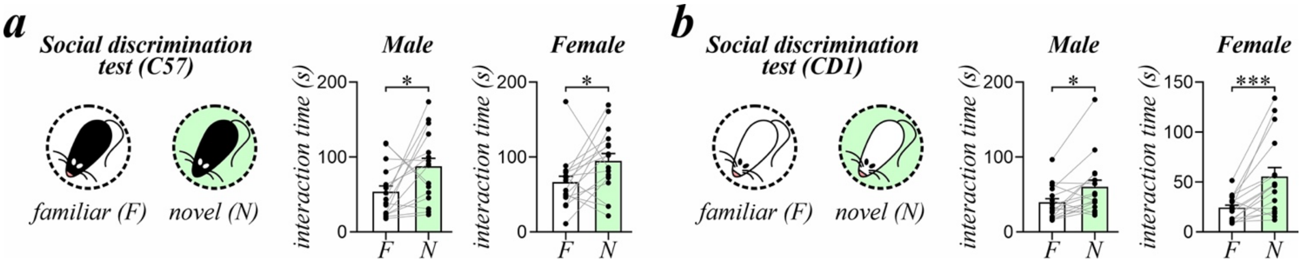


**Fig. S2. Social novelty recognition in male and female mice.**
Interaction time during the social discrimination test using familiar (F, white) and novel (N, green) (**A**) C57BL/6 and (**B**) CD1 mice. Data presented as mean ± SEM. *$p < 0.05$ ***$p < 0.001$, Wilcoxon test.

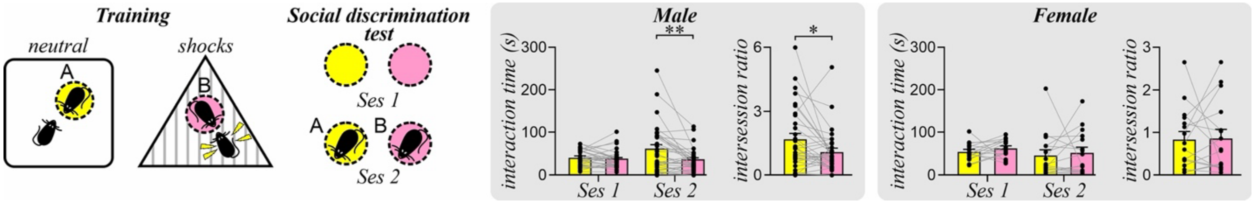


**Fig. S3. Sex difference in identity recognition in the negative social valence (NSV) tests with C57BL/6 targets.**

Behavioral schematic of the NSV task using C57BL/6 targets. Neutral trials with mouse A (yellow) and NSV trials with mouse B (pink). Histograms show the interaction time during session 1 (Ses 1, empty cups) and session 2 (Ses 2, with conspecific mice) of the social discrimination test, and the intersession ratio (Ses 2/Ses 1 interaction time) for each conspecific. Data presented as mean ± SEM. *p < 0.05, **p < 0.01, Wilcoxon test with Bonferroni correction.

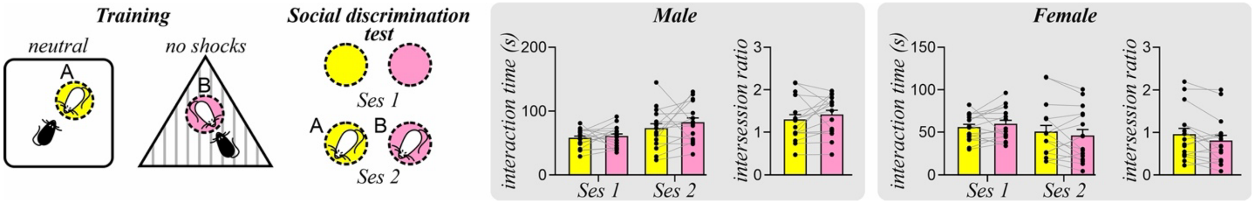


**Fig. S4. Identity recognition with no shock association.**
Behavioral schematic of the NSV task with no delivered shocks. Neutral trials with mouse A (yellow) and no-shock trials with mouse B (pink). Histograms show the interaction time during session 1 (Ses 1, empty cups) and session 2 (Ses 2, with conspecific mice) of the social discrimination test, and the intersession ratio (Ses 2/Ses 1 interaction time) for each conspecific. Data presented as mean ± SEM.

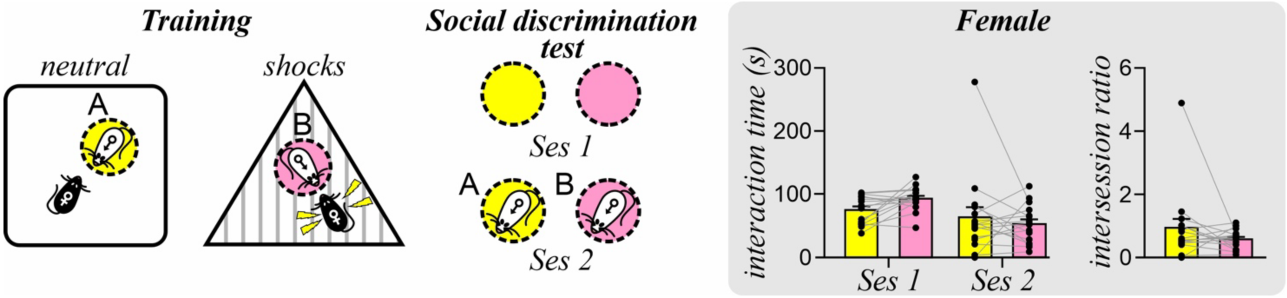


**Fig. S5. Identity recognition in female mice after the negative social valence (NSV) task with opposite sex targets.**

Behavioral schematic of the NSV task with opposite sex conspecific mice. Neutral trials with male mouse A (yellow) and NSV trials with male mouse B (pink). Histograms show the interaction time during session 1 (Ses 1, empty cups) and session 2 (Ses 2, with conspecific mice) of the social discrimination test, and the intersession ratio (Ses 2/Ses 1 interaction time) for each conspecific. Data presented as mean ± SEM.

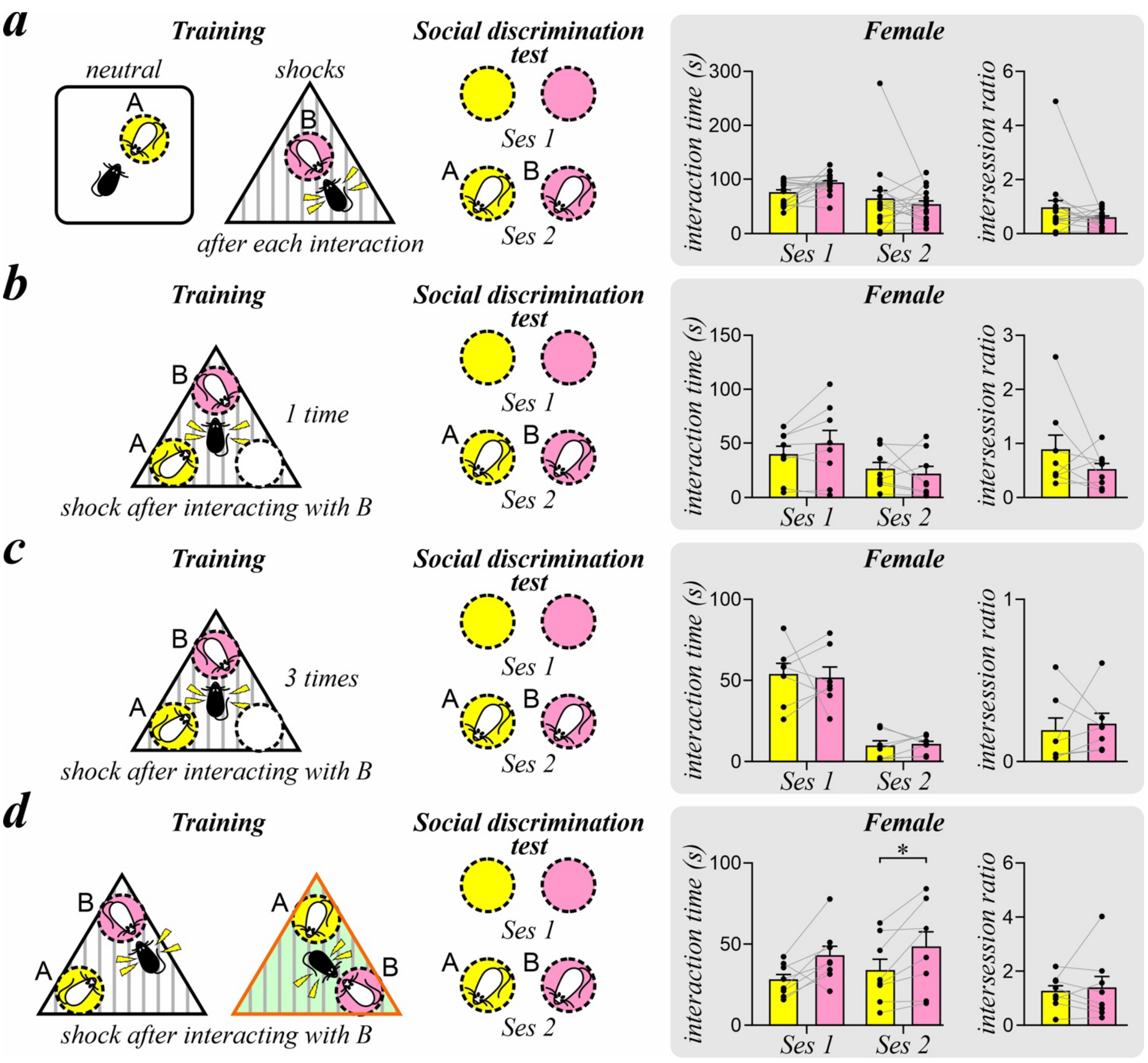


**Fig. S6. Identity recognition in female mice after variations of the (negative social valence) NSV task.**
Behavioral schematics of the NSV task in different variations. All variations depict neutral experiences with mouse A (yellow) and NSV experiences with mouse B (pink). Histograms show the interaction time during session 1 (Ses 1, empty cups) and session 2 (Ses 2, with conspecific mice) of the SDT, and the intersession ratio (Ses 2/Ses 1 interaction time) for each conspecific.
(**A**) Foot shocks delivered after each interaction bout in the NSV trial with mouse B.
(**B**) Training took place in a shock box containing three cups: empty (white), with mouse A, and with mouse B. Foot shocks were only delivered after each interaction bout with mouse B.
(**C**) Training was similar to (**B**) but repeated 3 times and with the cup locations rotated in each trial. (**D**) Two training sessions were performed in 2 different contexts. Foot shocks were only delivered after each interaction bout with mouse B. Data presented as mean ± SEM. $*p < 0.05$; Wilcoxon test with Bonferroni correction.

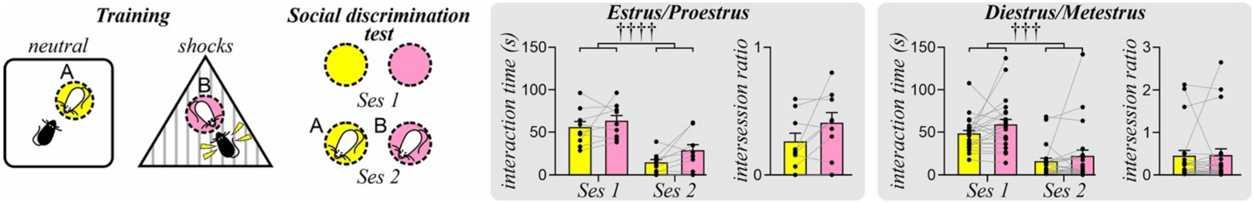


**Fig. S7. Identity recognition after the negative social valence (NSV) task does not vary with estrous cycle stage in female mice.**

Behavioral schematic of the NSV task with estrous cycle monitoring. Neutral trials with mouse A (yellow) and NSV trials with mouse B (pink). Histograms show the interaction time during session 1 (Ses 1, empty cups) and session 2 (Ses 2, with conspecific mice) of the social discrimination test, and the intersession ratio (Ses 2/Ses 1 interaction time) for each conspecific. At the time of the SDT subjects were in proestrus/estrus or metestrus/diestrus stages. Data presented as mean ± SEM. ††††$p < 0.0001$, †††$p < 0.001$, effect test of session with two-way ANOVA.

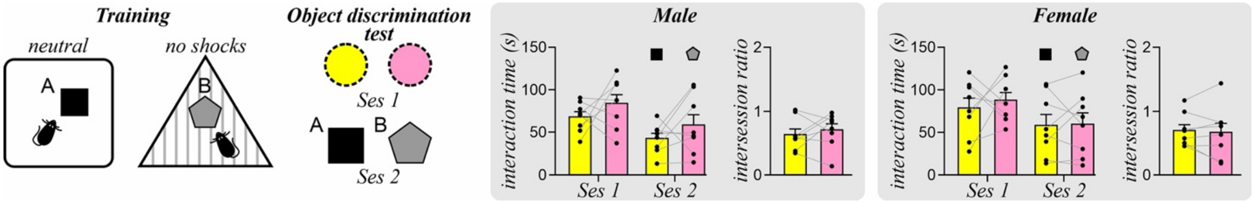


**Fig. S8. Object recognition with no emotional valence association.**
Behavioral schematic of the task. Neutral trials with object A (yellow) and object B (pink) in 2 different contexts. Histograms show the interaction time during session 1 (Ses 1, empty cups) and session 2 (Ses 2, with objects) of the discrimination test, and the intersession ratio (Ses 2/Ses 1 interaction time) for each object. Data presented as mean ± SEM.

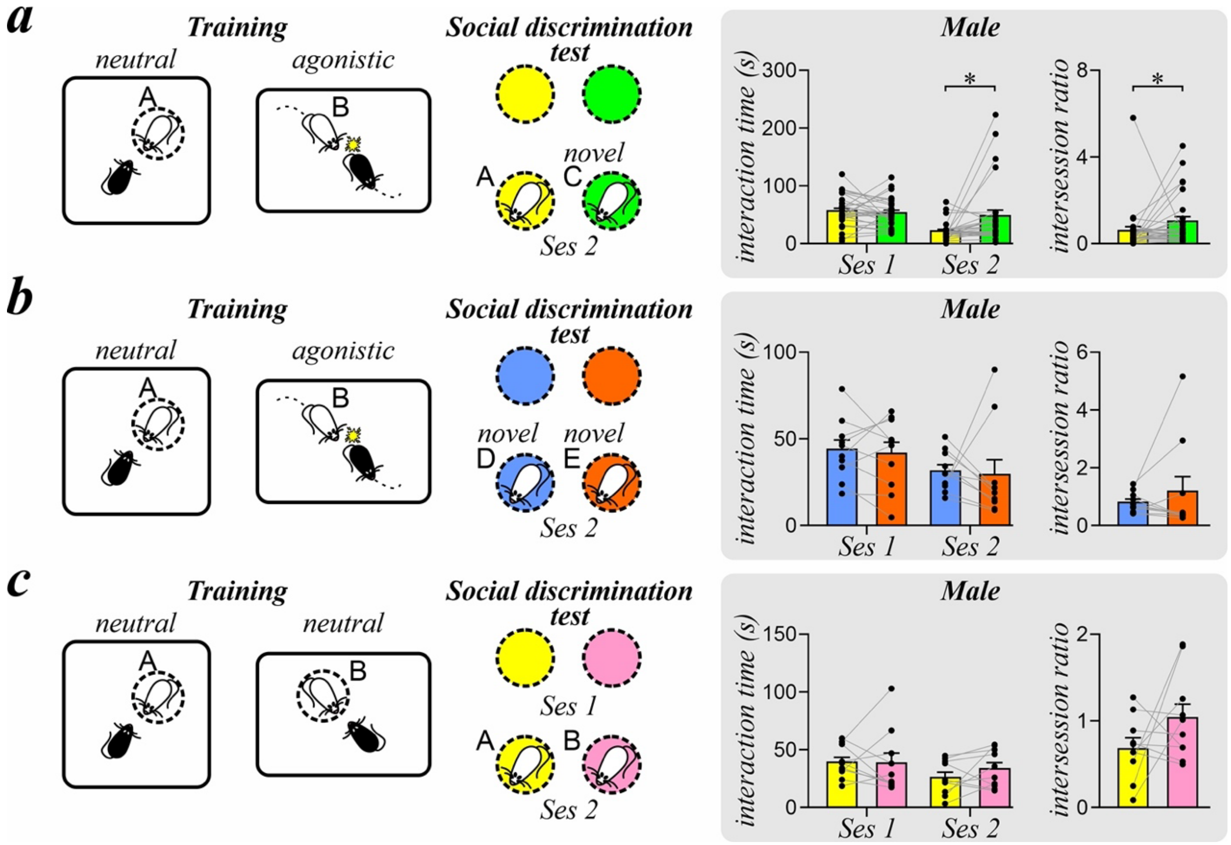


**Fig. S9. Identity recognition in male mice after variations of the negative social valence-social defeat stress (NSV-SDS) task.**

Behavioral schematics of the NSV-SDS task in different variations. Neutral trials with mouse A (yellow) and NSV-SDS trials with mouse B (pink). Histograms show the interaction time during session 1 (Ses 1, empty cups) and session 2 (Ses 2, with conspecific mice) of the social discrimination test, and the intersession ratio (Ses 2/Ses 1 interaction time) for each conspecific. (**A**) SDT with neutral mouse A (yellow) and novel mouse C (green). (**B**) SDT with two novel mice D (blue) and E (red). (**C**) No-attack variation where both mice were confined within enclosures. Data presented as mean ± SEM. $*p < 0.05$, Wilcoxon test with Bonferroni correction.

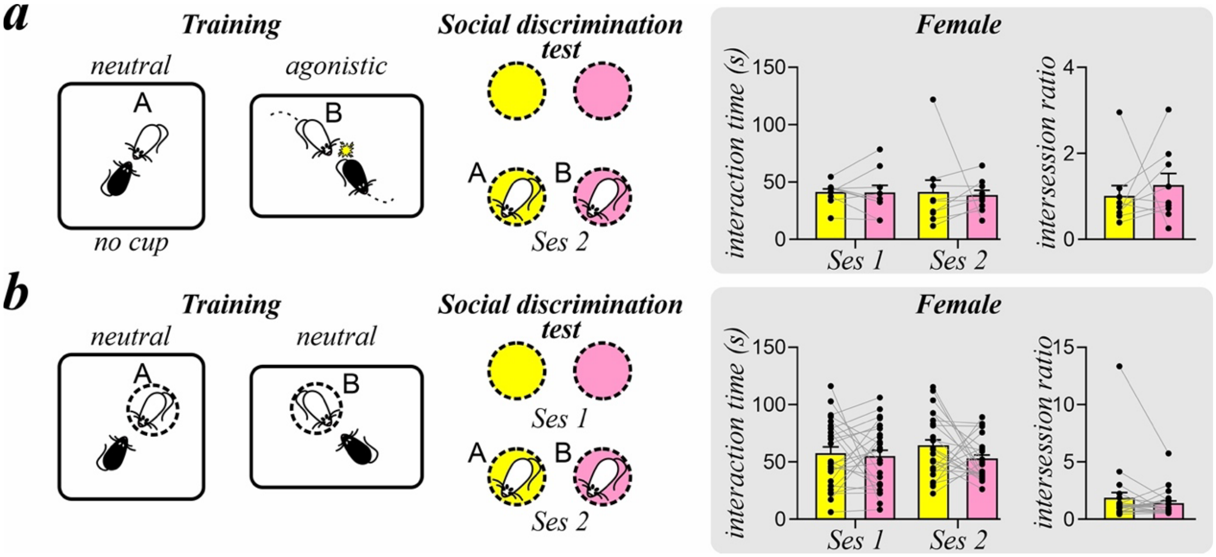


**Fig. S10. Identity recognition in female mice after variations of the negative social valence-social defeat stress (NSV-SDS) task.**

Behavioral schematics of the NSV-SDS task in different variations. Neutral trials with mouse A (yellow) and NSV-SDS trials with mouse B (pink). Histograms show the interaction time during session 1 (Ses 1, empty cups) and session 2 (Ses 2, with conspecific mice) of the social discrimination test, and the intersession ratio (Ses 2/Ses 1 interaction time) for each conspecific. (**A**) Neutral mice A was non-aggressive, allowing for both conspecifics to be freely moving. (**B**) No-attack variation where both conspecifics were confined within enclosures. Data presented as mean ± SEM.

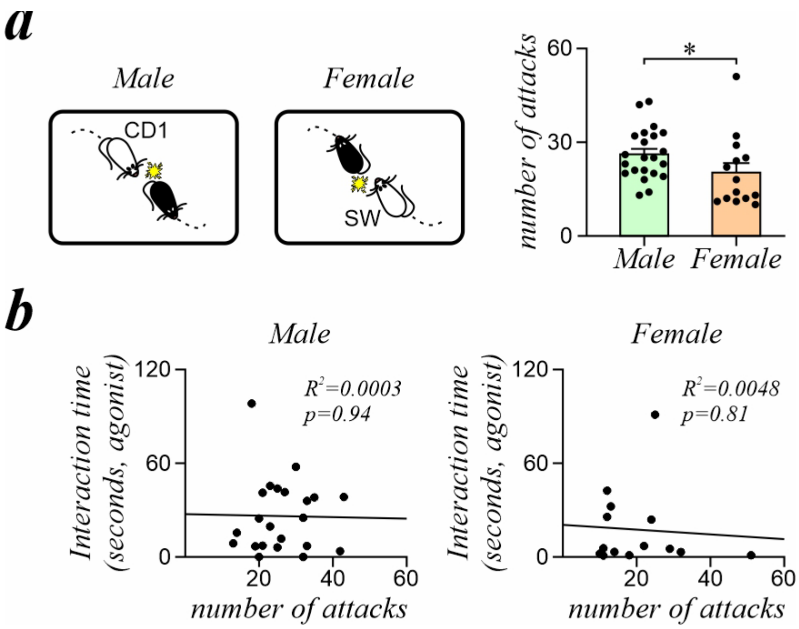


**Fig. S11: Attacks by aggressors in negative social valence-social defeat stress (NSV-SDS) tasks in male and female mice.**

(**A**) *Left:* Male and female C57BL/6 mice were attacked by male CD1 and female SW mice during NSV-SDS, respectively. *Right:* Histograms show the number of attacks by male or female aggressors during the NSV-SDS trial. (**B**) Scatter plots show the relationship between the number of attacks from aggressors and the interaction time with these aggressors during the social interaction test in male and female mice. Data presented as mean ± SEM. $^*p<0.05$, Mann Whitney test.

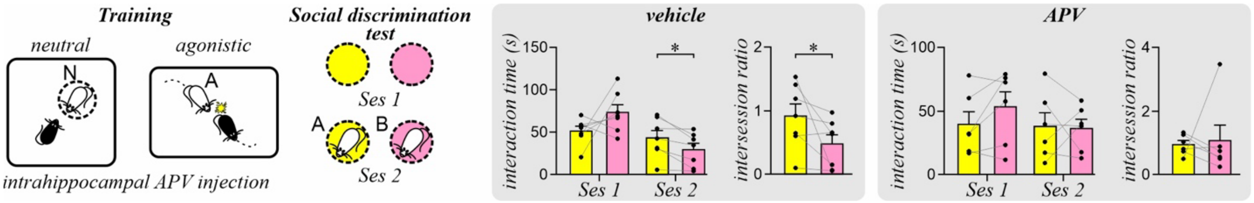


**Fig. S12: Inhibition of dCA1 activity abolishes identity recognition in male mice after the negative social valence-social defeat stress (NSV-SDS) task.**

Behavioral schematics of the NSV-SDS task. Bilateral intrahippocampal APV or vehicle injections were administered prior to each trial. Neutral trials with mouse A (yellow) and NSV-SDS trials with mouse B (pink). Histograms show the interaction time during session 1 (Ses 1, empty cups) and session 2 (Ses 2, with conspecific mice) of the SDT, and the intersession ratio (Ses 2/Ses 1 interaction time) for each conspecific. Data presented as mean ± SEM. *$p < 0.05$, Wilcoxon test with Bonferroni correction.

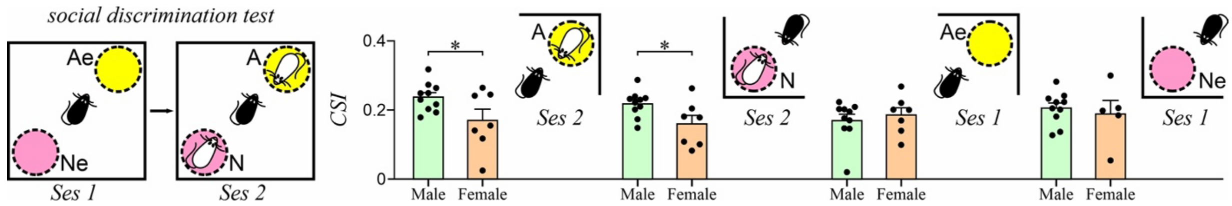


**Fig. S13: Cosine similarity index of social and nonsocial ensembles in male and female mice.**
*Left:* Schematic diagram of the SDT after NSV-SDS. *Right:* Histograms show the cosine similarity index (CSI) of social (for mouse-containing wire cups A or N) and nonsocial ensembles (empty wire cups Ae or Ne) in male and female mice. Data presented as mean ± SEM. *$p<0.05$, Student's t-test.


1. E. L. Newman *et al.*, Fighting Females: Neural and Behavioral Consequences of Social Defeat Stress in Female Mice. *Biol Psychiatry* **86**, 657-668 (2019).
2. S. S. Moy *et al.*, Sociability and preference for social novelty in five inbred strains: an approach to assess autistic-like behavior in mice. *Genes Brain Behav* **3**, 287-302 (2004).
3. C. S. Caligioni, Assessing reproductive status/stages in mice. *Curr Protoc Neurosci* **Appendix 4**, Appendix 4I (2009).
4. S. L. Resendez *et al.*, Visualization of cortical, subcortical and deep brain neural circuit dynamics during naturalistic mammalian behavior with head-mounted microscopes and chronically implanted lenses. *Nat Protoc* **11**, 566-597 (2016).
5. E. A. Pnevmatikakis, A. Giovannucci, NoRMCorre: An online algorithm for piecewise rigid motion correction of calcium imaging data. *J Neurosci Methods* **291**, 83-94 (2017).
6. P. Zhou *et al.*, Efficient and accurate extraction of in vivo calcium signals from microendoscopic video data. *Elife* **7**, (2018).
7. A. Mathis *et al.*, DeepLabCut: markerless pose estimation of user-defined body parts with deep learning. *Nat Neurosci* **21**, 1281-1289 (2018).
8. L. Carrillo-Reid, V. G. Lopez-Huerta, M. Garcia-Munoz, S. Theiss, G. W. Arbuthnott, Cell Assembly Signatures Defined by Short-Term Synaptic Plasticity in Cortical Networks. *Int J Neural Syst* **25**, 1550026 (2015).
9. J. P. Hamm, D. S. Peterka, J. A. Gogos, R. Yuste, Altered Cortical Ensembles in Mouse Models of Schizophrenia. *Neuron* **94**, 153-167 e158 (2017).
